# AI-Driven Insights into the complexities of Chinese hamster ovary cells death in order to optimize production processes

**DOI:** 10.1101/2023.11.14.567093

**Authors:** Mohammad Moshfeghnia

## Abstract

Chinese hamster ovary (CHO) cells are a multipurpose and high-performance cell line for recombinant protein production in biopharmaceutical industry. They have proven their ability to produce a wide range of therapeutic proteins with high efficiency and quality. Designing novel and high-performance CHO cell lines has an incredible impact in biopharmaceutical industry that can reduce prices and increase product efficiency. One of the best ways is to prevent CHO cells death during Bioprocessing. Apoptosis is the most common form of CHO cells death during Bioprocessing. Analyzing Apoptosis and cell-cycle complex signaling pathways are necessary for the control of cell growth, efficiency, and the death of CHO cells. Therefore, analyzing and understanding interactions of these pathways and their interactions with other cellular processes can help optimize the performance and quality of CHO cell lines. AI-driven insight solutions and Advanced machine learning algorithms like GAT (Graph Attention Network) used in this project indicate most important Targets in complex signaling pathways. Pathways such as the TNF signaling pathway, and also viruses like: Hepatitis C, HIV1 and Bacteria like: Salmonella have High intersection size and Low P-value with complex signaling pathways. These microorganisms should be used to design high-performance CHO cell lines because they are master in these pathways. This method can be used to find novel and high efficiency targets for curing cancer in humans.

## Introduction

Chinese hamster ovary (CHO) cells are the most widely used mammalian cell line for the production of recombinant protein therapeutics, especially monoclonal antibodies (mAbs). CHO cells have several advantages over other cell lines, such as their ability to grow in suspension culture at high densities, their adaptability to various growth conditions, and their capacity to perform proper folding and post-translational modifications of the expressed proteins [1].

CHO cells have a long history of use in biopharmaceutical industry, dating back to 1987 when the first recombinant therapeutic protein expressed in CHO cells, tissue plasminogen activator (tPA), was approved for clinical use [1]. Since then, many other recombinant proteins have been successfully produced in CHO cells, such as erythropoietin, interferons, growth factors, coagulation factors, and hormones [1]. However, the most prominent application of CHO cells is in the production of mAbs, which account for more than half of all biopharmaceuticals on the market [3]. CHO cells are able to produce mAbs with high yield and quality, as well as with human-like glycosylation patterns [4].

The development of a CHO cell line for recombinant protein production involves several steps, such as transfection of the gene encoding the protein of interest into a host cell line, selection and screening of expressing clones, characterization and optimization of cell culture parameters, scale-up and validation of the production process, and analysis of the product quality [2]. Each step requires careful consideration and optimization to achieve the desired productivity and quality of the recombinant protein. Moreover, each step may introduce genetic and phenotypic variations in the CHO cell line, which may affect its performance and stability over time [2]. Therefore, it is important to monitor and control the genetic and environmental factors that influence the CHO cell line during its development and manufacturing.

CHO cells are not only a valuable tool for biopharmaceutical production, but also a fascinating model system for studying cellular biology and physiology. The genome sequence of CHO cells was published in 2011, revealing insights into their genomic features and evolution [5]. Since then, various omics technologies have been applied to study the transcriptome, proteome, metabolome,and glycome of CHO cells, providing a comprehensive understanding of their molecular mechanisms and pathways [6]. These studies have also enabled the identification of key genes and factors that regulate the expression and glycosylation of recombinant proteins in CHO cells [6]. Furthermore, gene editing technologies such as CRISPR/Cas9 have been used to manipulate the genome of CHO cells to improve their characteristics and performance as biopharmaceutical hosts [7].

CHO cells have several advantages, such as high growth rate, adaptability to various culture conditions, and ability to perform complex post-translational modifications, especially glycosylation [2]. The biopharmaceutical industry is constantly seeking to improve the productivity and quality of CHO-derived products, as well as to reduce the cost and time of development and manufacturing.

The global market for biopharmaceuticals produced using CHO cells is expected to grow steadily in the next few years, driven by the increasing demand for monoclonal antibodies, biosimilars, and novel biologics for various indications. According to Global Data, the top 20 biopharmaceutical companies reported a marginal increase in their market capitalization from $3.56 trillion in Q2 2023 to $3.57 trillion in Q3 2023 [8].

CHO cell lines are widely used for the production of recombinant proteins and monoclonal antibodies in biopharmaceutical industry [9]. However, the productivity and quality of these products are often limited by the intrinsic characteristics of CHO cells, such as low specific productivity, inefficient protein folding and secretion, and susceptibility to apoptosis [10]. Therefore, there is a great need to design novel and high-performance CHO cell lines with genetic and protein engineering.

Genetic engineering can introduce or modify genes that enhance the expression, folding, secretion, or stability of the target proteins [11]. Protein engineering can modify the structure and function of the target proteins to improve their affinity, specificity, or activity [12]. By combining these two approaches, it is possible to create CHO cell lines that can produce high-quality and high-value biopharmaceuticals with improved efficacy and safety [13].

One of the challenges in biotechnology is to produce high performance CHO cell lines that can withstand various stress conditions and maintain high productivity [9]. Apoptosis, or programmed cell death, is a major factor that affects the viability and productivity of CHO cells [14]. Apoptosis can be triggered by two main pathways: **1. extrinsic pathway** and **2. intrinsic pathway** [15]. The extrinsic pathway involves the activation of death receptors on the cell surface, such as Fas, TNFR1, DR3, DR4, and DR5, which belong to the TNF (Tumor Necrosis Factor) gene superfamily [16]. These receptors recruit different signaling molecules that activate caspase-8, a key initiator of apoptosis [17]. The intrinsic pathway is activated by intracellular stress signals, such as growth factor withdrawal, toxin accumulation, or Ca2+ flux, that cause mitochondrial dysfunction and release of pro-apoptotic proteins, such as cytochrome c and Smac/DIABLO [18]. These proteins activate caspase-9, another initiator of apoptosis [19]. Both caspase-8 and caspase-9 then cleave and activate effector caspases, such as caspase-3, -6, and -7, which execute apoptosis by cleaving various cellular substrates [20].

Understanding the molecular mechanisms of apoptosis in CHO cells is crucial for developing strategies to enhance their performance and productivity. By modulating the expression or activity of death receptors, signaling molecules, or caspases, it may be possible to increase the resistance of CHO cells to apoptosis and improve their survival and yield. Therefore, we should focus on death signaling pathways, especially apoptosis, to produce high performance CHO cell lines [21].

Apoptosis can reduce cell viability, productivity, and product quality, as well as increase the complexity and cost of downstream processing. Therefore, controlling apoptosis is essential for designing high performance CHO cell lines for biopharmaceutical production [21].

Apoptosis can be triggered by various stresses that cells encounter during bioprocessing, such as hypoxia, shear, nutrient depletion, pH, osmolality, and protein misfolding. These stresses can activate different apoptotic pathways, involving caspases, Bcl-2 family proteins, and other molecules, that lead to cell death. Understanding the mechanisms and regulation of apoptosis in CHO cells can help to identify the key factors that influence cell survival and function [21].

Several strategies have been proposed to control apoptosis in CHO cell lines, such as optimizing culture media and operating conditions, engineering cell lines to overexpress anti-apoptotic genes or under express pro-apoptotic genes, and developing mathematical models to simulate and optimize bioprocesses. These strategies aim to minimize the apoptotic stresses, enhance the cell resistance, and improve the culture performance [21].

Controlling apoptosis is important not only for increasing the quantity, but also the quality of the bioproducts. Apoptosis can affect the glycosylation, folding, and aggregation of recombinant proteins, which can have implications for their efficacy, safety, and immunogenicity. Therefore, controlling apoptosis can ensure the consistency and quality of the bioproducts, as well as facilitate the quality-by-design approach in bioprocess development [21].

In conclusion, apoptosis is a critical factor that determines the performance of CHO cell lines in bioprocessing. Controlling apoptosis can improve cell viability, productivity, and product quality, as well as reduce the bioprocess complexity and cost. Therefore, controlling apoptosis is essential for designing high performance CHO cell lines for biopharmaceutical production [21].

Cell-cycle signaling pathways regulate the growth, proliferation and differentiation of cells, while apoptosis signaling pathways control the programmed cell death or elimination of unwanted cells [22].

Both types of pathways are important for maintaining the homeostasis and quality of CHO cell lines, which are widely used for the production of recombinant proteins and biopharmaceuticals [9]. By analyzing both cell-cycle and apoptosis signaling pathways, we can gain a better understanding of the molecular mechanisms that affect the productivity, stability and robustness of CHO cell lines [10]. This can lead to the development of more efficient and cost-effective bioprocesses for the manufacturing of therapeutic proteins and other bioproducts [13].

Apoptosis and cell-cycle signaling pathways are crucial for the regulation of cell growth, differentiation, and death in mammalian cells [15]. These pathways are also involved in the production of recombinant proteins using CHO cells, which are the most widely used host cells for biopharmaceutical manufacturing [10]. Therefore, understanding the molecular mechanisms of these pathways and their interactions with other cellular processes can help to optimize the performance and quality of CHO cell lines.

Advanced machine learning algorithms and AI solutions can provide powerful tools to analyze large-scale data sets from omics technologies like proteomics that analyze apoptosis and cell-cycle signaling pathways [23]. By applying these methods, we can gain insights into the complex regulatory networks that govern these pathways and identify key factors that influence their activity and stability. Moreover, we can use these insights to design high-performance CHO cell lines that have enhanced productivity, robustness, and consistency [24].

Machine learning algorithms, especially graph neural networks (GNNs), have shown great potential for analyzing complex biological networks, such as apoptosis and cell-cycle protein-protein interactions (PPIs) [25]. These networks are crucial for understanding the molecular mechanisms of cell growth, survival, and death, which are relevant for designing high-performance CHO cell lines for recombinant protein production [26]. In this paper, we propose a GNN-based model that can analyze PPIs from graph structure information. Our model leverages the advantages of representation learning and graph-based deep learning to capture the features and patterns of proteins and their interactions. We show that our model achieves superior performance and can provide insights into the biological roles of proteins and their interactions. This GNN-based model can also be used to recognize most important targets that lead cancer in humans. Actually, we can experiment with our targets in CHO cell line to find novel bioengineering methods for cure cancer in humans.

## Methods

Graph neural networks (GNNs) are a powerful machine learning technique that can learn from protein-protein interactions (PPIs). GNNs can encode the features of each protein node and its neighbors in a graph, and use graph convolutions to aggregate and propagate information across the network. This way, GNNs can capture the complex and dynamic patterns of PPIs interactions [25] [27]. Traditional machine learning algorithms, such as support vector machines (SVMs) or random forests (RFs), rely on hand-crafted features or sequence-based embeddings to represent proteins. However, these methods often ignore the structural information of PPI networks, such as their degree, position, and neighboring nodes in a graph [25] [28].

Therefore, we argue that GNNs are a better choice for analyzing PPIs interactions. GNNs can learn unknown from correlations, that is, they can infer the interactions of novel proteins based on their similarities and connections with known proteins. GNNs can also handle different types and scales of PPI networks, such as apoptosis and cell-cycle signaling pathways, by adapting their graph convolutions [27].We recommend using GNNs instead of traditional machine learning algorithms for PPI analysis.

Graph Neural Networks (GNNs) are a class of artificial neural networks that can process data represented as graphs. Graphs are data structures that consist of nodes (or vertices) and edges (or links) that connect them. Nodes can represent any entity and edges can represent any relationship. Graphs are useful for modeling complex systems with interactions and dependencies [28].

There are many types of GNNs, but they all share some common characteristics. They all use a process called message passing, which means that each node updates its representation (or feature vector) by aggregating information from its neighboring nodes. This way, nodes can learn from their local context and capture the structure of the graph [28].

One of the most High-performance types of GNNs is the Graph Attention Network (GAT), which was proposed by Velickovic et al. in 2018 [29]. GAT is based on the idea of self-attention, which means that each node learns how to assign different importance weights to its neighbors. This way, GAT can focus on the most relevant nodes for each node’s representation, and also handle graphs with varying degrees of nodes. GAT uses a multi-head attention mechanism, which means that each node computes multiple attention coefficients for each neighbor, and then concatenates or averages them to form the final node embedding. GAT has been shown to outperform other GNNs on various tasks, such as node classification [29]. One of the potential applications of GAT is in bioinformatics, where graphs can be used to model biological pathways. For example, GAT can be used to analyze apoptosis and cell-cycle signaling pathways, which are networks of proteins that regulate cell death. By applying GAT to these pathways, we can identify the most important proteins (or targets) that influence the outcome of these processes. This can help us understand the mechanisms of apoptosis and cell-cycle to control them.

A Graph Attention Network (GAT) is a machine learning algorithm that operates on graph-structured data. It uses masked self-attentional layers to learn different weights for different nodes in a neighborhood, without requiring any matrix operations or prior knowledge of the graph structure [29]. The architecture of a GAT consists of stacking multiple attention layers, where each layer computes a linear transformation of the node features, followed by a softmax function to obtain the attention coefficients. The output of each layer is then a weighted sum of the transformed features of the neighboring nodes [29]. We use GAT machine learning algorithm for first time to find most important protein targets for design novel CHO cell lines that cause optimize production processes and reduce costs. Our GAT algorithm is type of Node classification due to the attention weights of each edge.

## GAT ARCHITECTURE

1. The input to our layer is a set of node features, 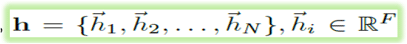, where *N* is the number of nodes, and *F* is the number of features in each node. The layer produces a new set of node features (of potentially different cardinality *F*′), 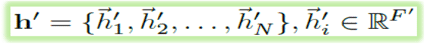, as its output [29].
2. In order to obtain sufficient expressive power to transform the input features into higher-level features, at least one learnable linear transformation is required. To that end, as an initial step, a shared linear transformation, parametrized by a weight matrix, **W** ∈ ℝ^*F*′ × *F*^, is applied to every node. We then perform self-attention on the nodes—a shared attentional mechanism computes attention coefficients *a* : ℝ^*F*′^ × ℝ^*F*′^ → ℝ. 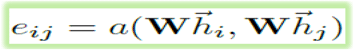 that indicate the importance of node *j*’s features to node *i*. The model allows every node to attend on every other node, dropping all structural information. We inject the graph structure into the mechanism by performing masked attention—we only compute *e*_*ij*_ for nodes *j* ∈ 𝒩_*i*_ where 𝒩_*i*_ is some neighborhood of node *i* in the graph. To make coefficients easily comparable across different nodes, we normalize them across all choices of *j* using the softmax function [29]:

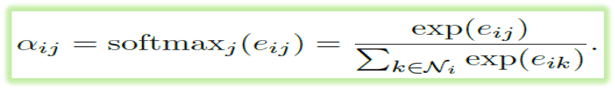
3. The attention mechanism *a* is a single-layer feedforward neural network, parametrized by a weight vector 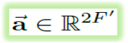, and applying the **LeakyReLU** nonlinearity. The coefficients computed by the attention mechanism may then be expressed as [29]:

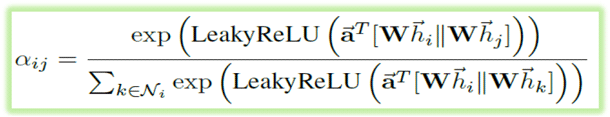 Where .*T* represents transposition and ‖ is the concatenation operation.
4. The normalized attention coefficients are used to compute a linear combination of the features corresponding to them, to serve as the final output features for every node (after potentially applying a nonlinearity, *σ*) [29]:

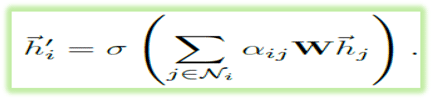
5. Multi-head attention mechanism: K independent attention mechanisms execute the transformation then their features are concatenated, resulting in the following output feature representation [29]:

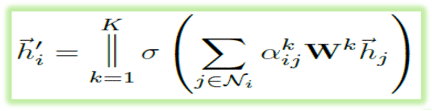 Where ‖ represents concatenation, 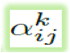 are normalized attention coefficients computed by the *k*-th attention mechanism (*a*^*k*^), and **W**^*k*^ is the corresponding input linear transformation’s weight matrix. The final returned output, **h**′ will consist of *KF*′ features (rather than *F*′) for each node.
6. we employ averaging, and delay applying the final nonlinearity [29]:

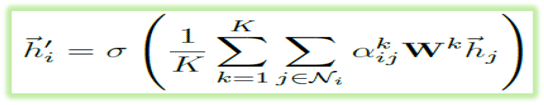

Main components of our Algorithm:

1. Packages and Libraries: Pandas is a popular Python library for data analysis and manipulation, which provides high-performance data structures and operations for working with tabular series data [31]. Numpy is another Python library that supports multidimensional arrays and matrices, as well as mathematical functions for linear algebra, statistics, and Fourier transform [32]. Torch is a Python package that provides tensor computations and automatic differentiation for deep learning [33]. Torch-geometric is an extension library for PyTorch that implements geometric deep learning methods for graphs [34]. Sklearn is a Python module that offers various tools for machine learning, such as classification [35]. Networkx is a Python package that enables the creation, manipulation, and analysis of complex networks, such as biological networks [36].
2. Datasets, Machine and software: Softwares : Jetbrains Dataspell for writing and editing codes [37] / Cytoscape for visualize our Algorithm [38] / Sublime text editor for editing codes [39] / Anaconda platform for perform machine learning Algorithm [40]. Machine: Training of Algorithm on Nvidia GeForce RTX 3060 GPU. Datasets: String-db version 12.0 Apoptosis and Cell-cycle of CHO datasets [41] / KEGG Apoptosis and Cell-cycle pathways, Also use KEGG pathways for enrichment of Algorithm [42]/ Scientific articles about Apoptosis and Cell-cycle in CHO cells used for validation our algorithm, According to these Articles [43 to 64] our nodes ( proteins ) divided into three categories : 1.Anti ( Anti Apoptosis ) 2.Pro ( Pro Apoptotic ) 3.Unknown ( Need Experimental Validation ).
3. Main parts of Algorithm : Importing necessary libraries / load the dataset / get unique node names and their labels / initialize node features with zero arrays / compute edge features and aggregate them to node features / normalize the aggregated features by the node degree (number of edges per node) / Leave-one-out cross-validation / Step the learning rate scheduler / Print the type of the attn tensor / Convert attn to a floating point tensor / Store attention weights to each edge / Taking the mean of all attention weights.

We use Leave-one-out cross-validation (LOOCV) for evaluating the performance of our machine learning model by using each observation in the dataset as a test set once and the rest of the observations as a training set. LOOCV has several advantages over other methods of cross-validation, such as k-fold or validation-set methods. First, LOOCV provides a less biased estimate of the test mean squared error (MSE) because it uses almost all the available data for training, thus reducing the impact of randomness in the data partitioning [65]. Second, LOOCV tends to not overestimate the test MSE as much as other methods, especially when the dataset is small or when the model is complex [66]. Third, LOOCV can be useful when there is limited labeled data available, as it does not require a separate validation set and uses all the data points for both training and testing [67].

Also, g_scs Significance Threshold Method is use for functional enrichment analysis. The g_scs Significance Threshold Method is a method for correcting multiple testing errors in functional enrichment analysis [68]. It is based on the Strong Control of the Generalized Family-wise Error Rate (gFWER), which is the probability of finding at least one false positive result among all significant results [69]. The g_scs method adjusts the p-values of each functional term by taking into account the hierarchical structure of the term ontology and the correlation between terms [68]. One of the strengths of this method is that it provides a better balance between sensitivity and specificity than other methods such as Bonferroni correction or false discovery rate (FDR) [68]. It also avoids over-penalizing terms that are highly dependent on each other, such as parent-child terms in the Gene Ontology [68]. Another strength of this method is that it can handle both flat and ordered gene lists, which allows for more flexible and informative analysis of gene expression data [69].

## Results

Our results are divided into two parts: 1. Apoptosis signaling pathway, and 2. Cell-cycle signaling pathway.

### 1. Apoptosis signaling pathway Algorithm

(Precision: 87%).

We display nodes that play a role in the Apoptosis signaling pathway (Figure 3). Each node has interactions with other nodes based on the weight of the edges. If the relationship between two nodes is stronger, they will have a higher edge weight. In our hierarchical layout, if an edge has a higher weight, it will also have greater thickness and a higher color contrast. We further enrich this hierarchical layout with the TNF signaling pathway, Hepatitis C, and the HIV-1 virus (Figure 6).

**Figure 1.**
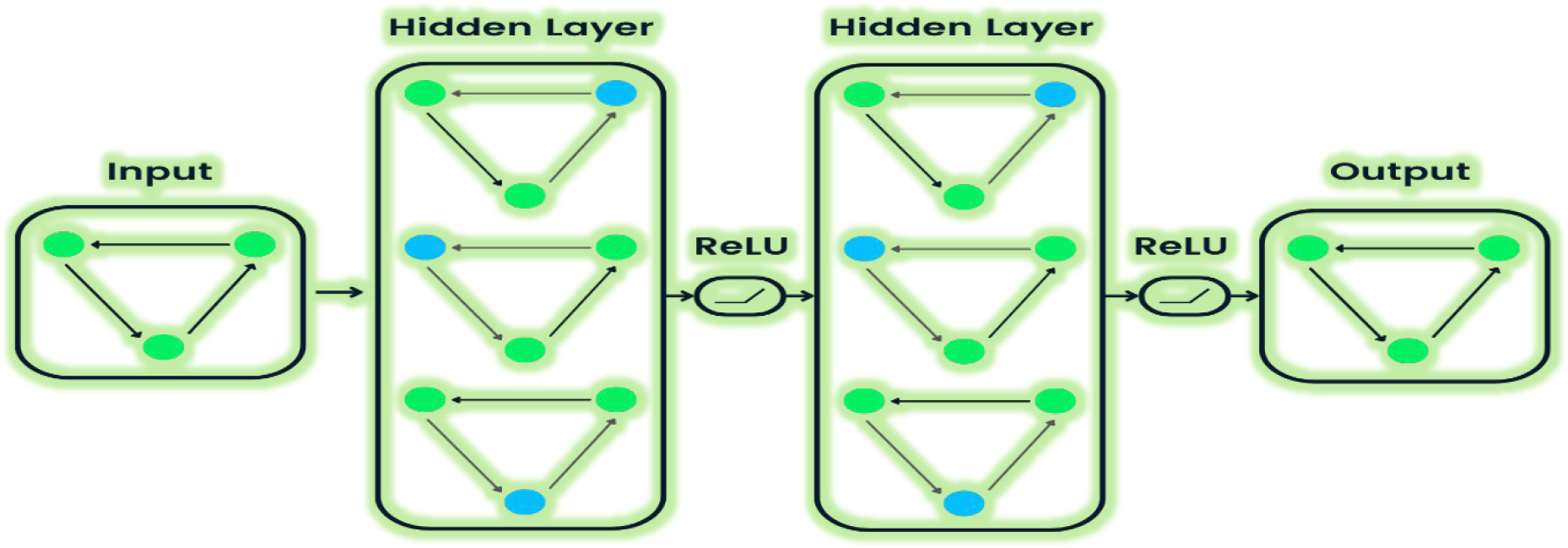
Basis of Graph Neural Networks (GNNs) Algorithms [30].

**Figure 2.**
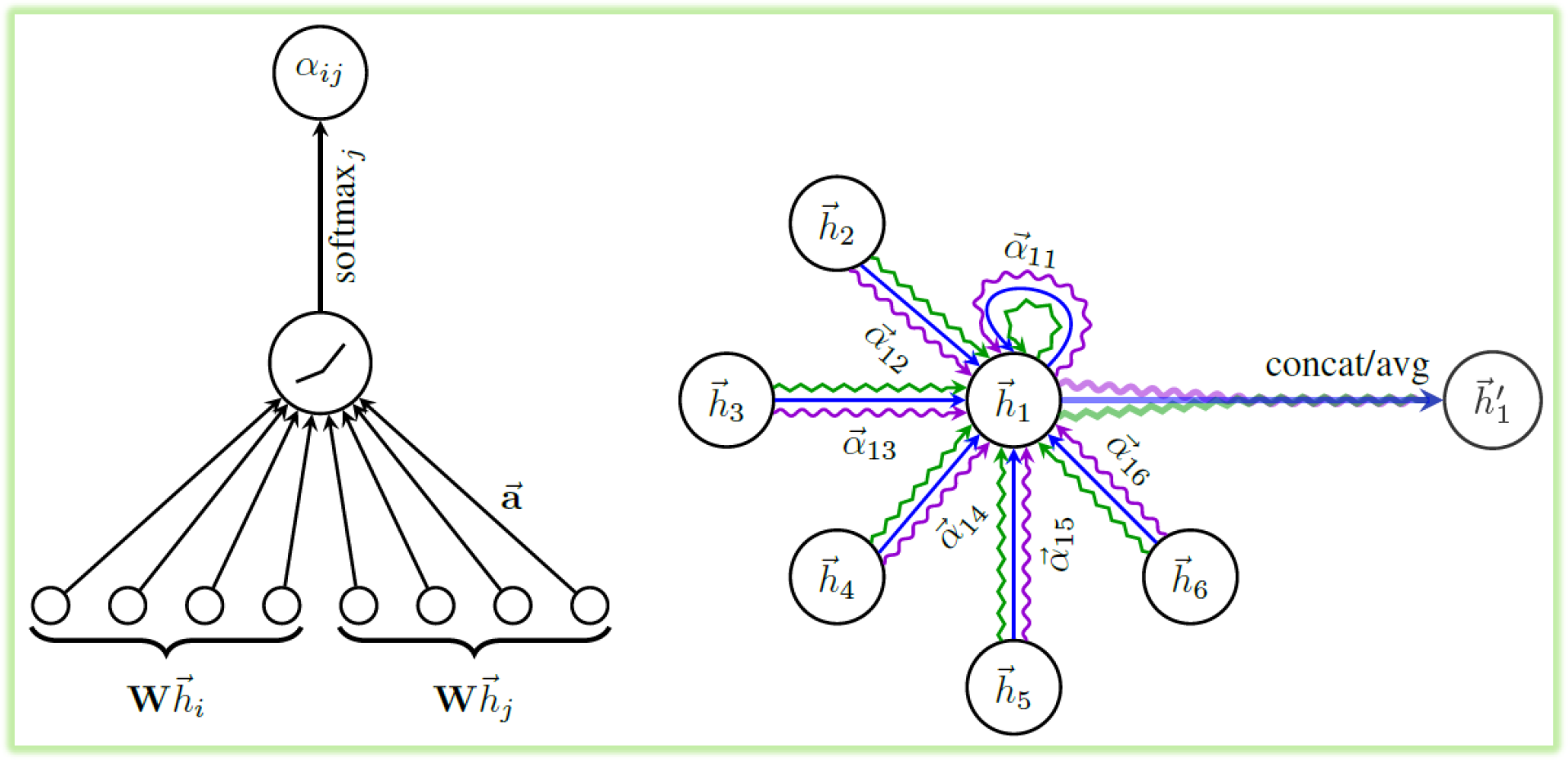
Left: The attention mechanism 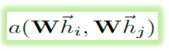 in our algorithm. Right: An illustration of multi head attention mechanism [29].

**Figure 3.**
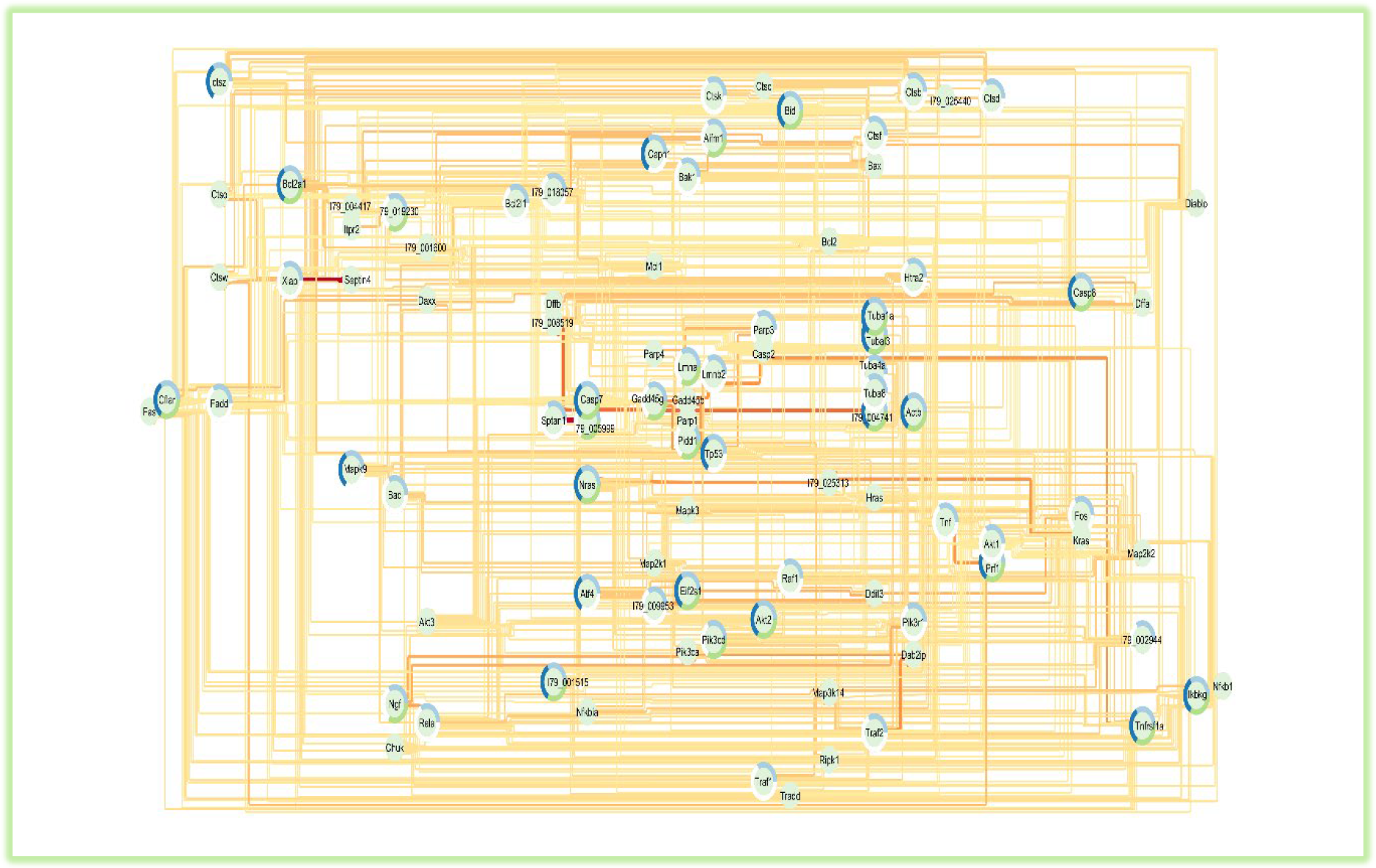
Hierarchic layout of GAT Apoptosis signaling pathway Algorithm.

Every node has important role in pathway, but we show the Twenty most important proteins in (Figure 4). This node ranking is based on criteria such as: 1. Stress, 2. Radiality, 3. Degree, and so on. Every node has one of these labels: 1. Anti (Anti-Apoptosis), 2. Pro (Pro-Apoptotic), 3. Unknown (Need Experimental Validation). Also, we show the twenty highest edge scores between nodes in (Figure 5).

**Figure 4.**
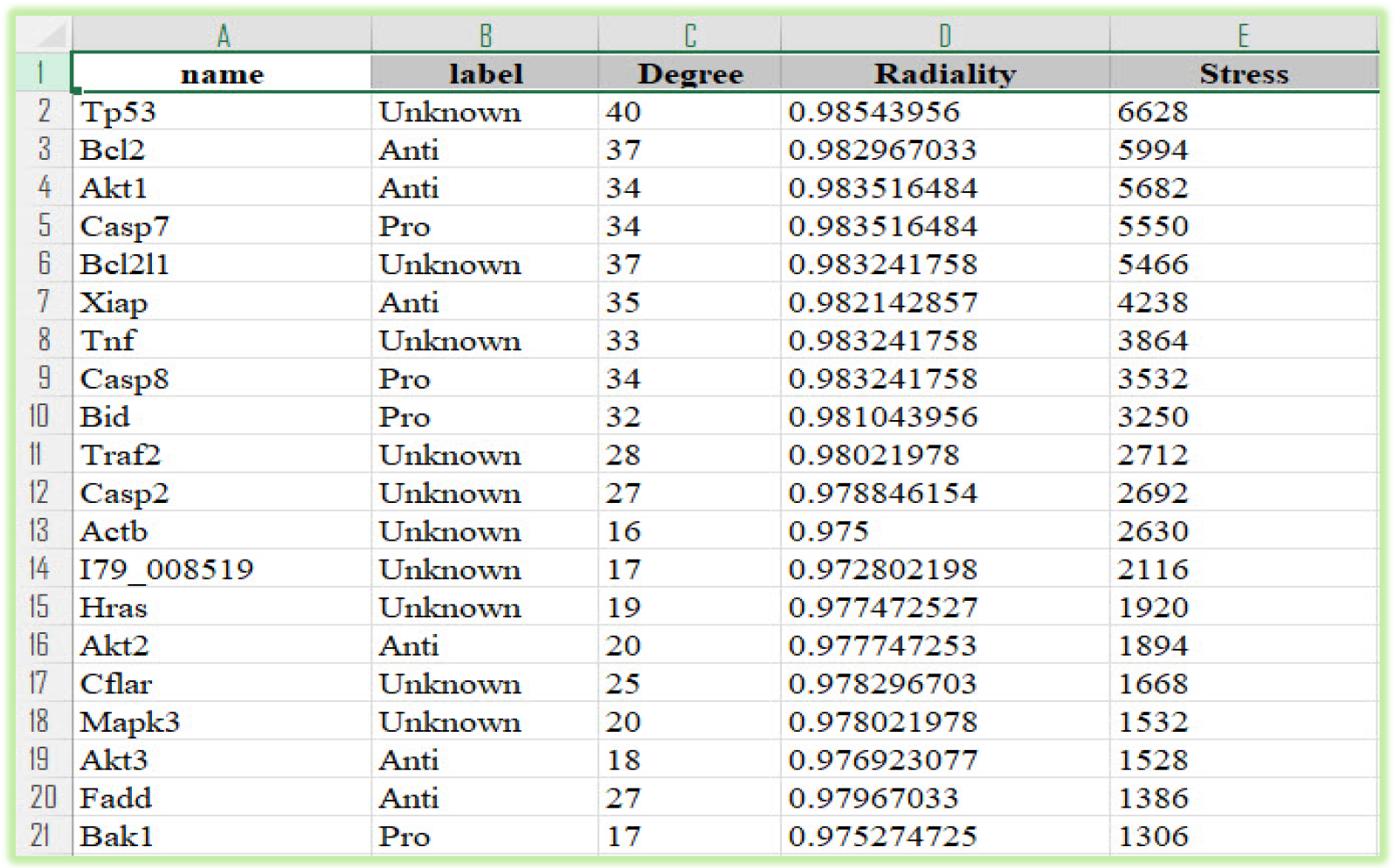
Most important nodes (proteins) in Apoptosis signaling pathway.

**Figure 5.**
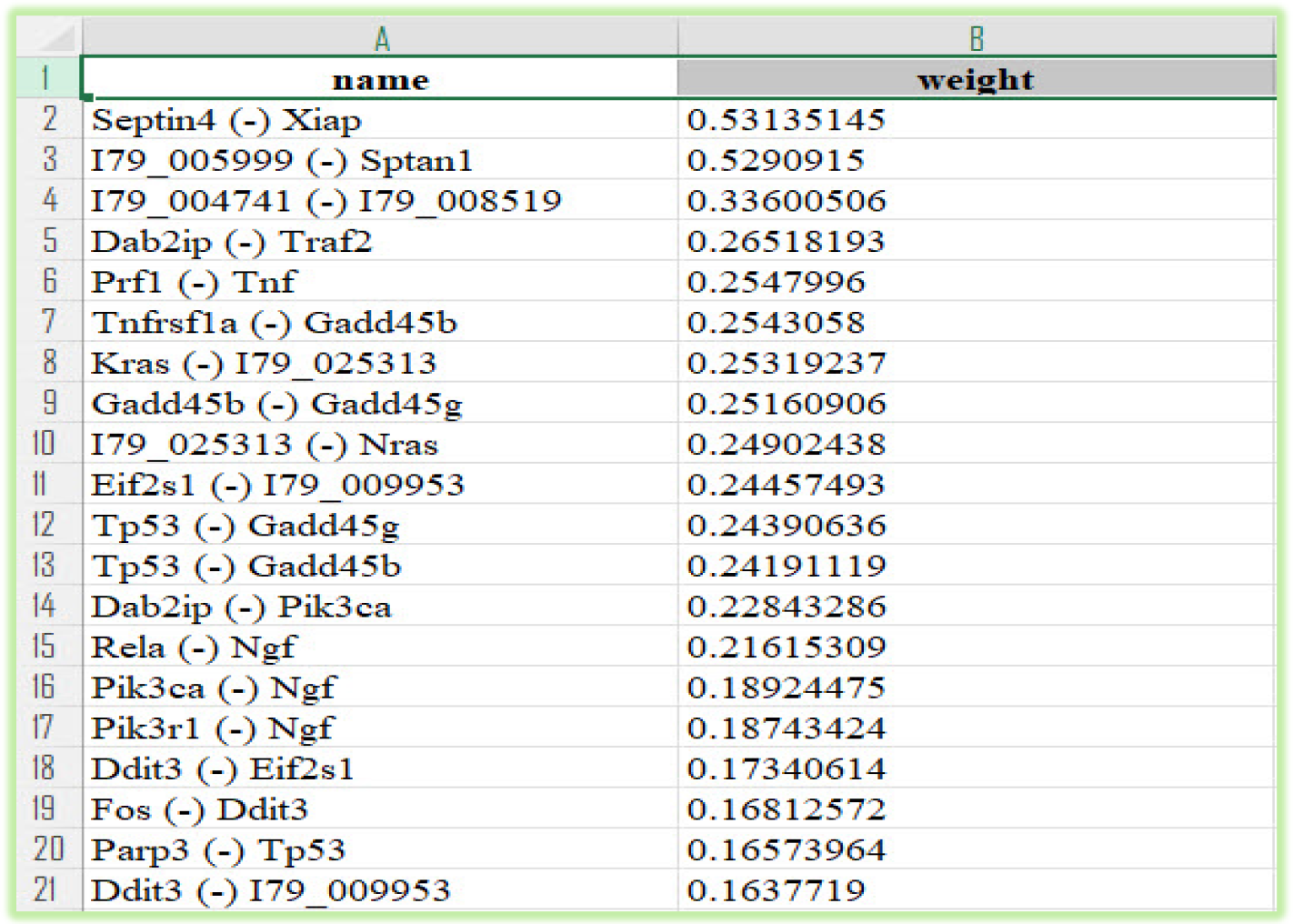
Most important Edges (Interactions between proteins) in Apoptosis signaling pathway.

#### 1.2. Most important nodes

1. **Tp53**: One of the most important nodes in Apoptosis signaling pathway is Tp53. Tp53 is a nuclear transcription factor that regulates the expression of various genes involved in apoptosis, growth arrest or senescence in response to genotoxic or cellular stress [71]. Tp53 is also known as the “guardian of the genome” because of its critical role in maintaining genomic integrity [72]. Tp53 is mutated in about half of all human cancers, which leads to loss of its tumor suppressive function and gain of oncogenic properties [72]. Tp53 mediates apoptosis through both extrinsic and intrinsic pathways, depending on the type and severity of the stress signal [71]. The extrinsic pathway of apoptosis is triggered by the binding of death ligands, such as FasL or TNF-alpha, to their respective receptors on the cell surface, which activate caspase-8 and initiate a cascade of caspase activation [71]. Tp53 can promote this pathway by upregulating the expression of death ligands and receptors, such as Fas, DR4, DR5 and PIDD [71]. Tp53 can also induce the expression of BH3-only proteins, such as Bid and Bim, which can translocate to the mitochondria and activate the intrinsic pathway [71]. The intrinsic pathway of apoptosis is initiated by the release of cytochrome c from the mitochondria into the cytosol, where it forms a complex with Apaf-1 and caspase-9, known as the apoptosome [71]. This complex activates caspase-9, which in turn activates caspase-3 and other downstream caspases [71]. Tp53 can induce this pathway by regulating the expression and activity of several members of the Bcl-2 family of proteins, which control the permeability of the mitochondrial outer membrane [71]. Tp53 can upregulate pro-apoptotic proteins, such as Bax, Bak, Puma and Noxa, and downregulate anti-apoptotic proteins, such as Bcl-2, Bcl-xL and Mcl-1 [71]. Tp53 can also modulate the activity of Bcl-2 family proteins by direct interaction or post-translational modifications [71]. The extrinsic and intrinsic pathways of apoptosis converge at the level of caspase-3 activation, which leads to the execution phase of apoptosis, characterized by DNA fragmentation, chromatin condensation, membrane blebbing and formation of apoptotic bodies [71]. Tp53 can also influence this phase by regulating the expression of genes involved in DNA repair, chromatin remodeling and phagocytosis [71]. In summary, Tp53 is a key regulator of apoptosis signaling pathways that can respond to different stress signals and induce cell death through various mechanisms. Tp53-mediated apoptosis is essential for preventing cancer development and progression.
2. **Bcl2**: Bcl2 is a protein that belongs to the Bcl-2 family of proteins that regulate cell death by either inhibiting or inducing apoptosis [104]. Apoptosis is a process of programmed cell death that is essential for maintaining tissue homeostasis and eliminating damaged or unwanted cells [105]. Bcl2 is an anti-apoptotic protein that prevents apoptosis by preserving the integrity of the mitochondrial membrane, which is a key site of apoptotic signaling [106]. Bcl2 also inhibits the activation of caspases, which are proteases that execute the apoptotic program [105]. Bcl2 is overexpressed in many hematological malignancies, such as chronic lymphocytic leukemia, follicular lymphoma and mantle cell lymphoma, where it confers resistance to chemotherapy and promotes tumor survival [107]. Therefore, Bcl2 is an important therapeutic target for these cancers, and several drugs that inhibit Bcl2, such as venetoclax, have been developed and approved for clinical use [107]. Bcl2 also plays a role in other diseases, such as hepatitis C virus infection, where it is upregulated and contributes to liver inflammation and fibrosis [108]. Bcl2 is a very important protein in apoptosis signaling pathway because it modulates the balance between cell survival and death, which has implications for various physiological and pathological processes. By inhibiting apoptosis, Bcl2 can protect cells from stress-induced damage, but also promote tumorigenesis and drug resistance. By targeting Bcl2, novel therapies can be designed to induce apoptosis in cancer cells and other diseases.
3. **Akt1**: Akt1 is a serine/threonine kinase that belongs to the PI3K/Akt signaling pathway, which is involved in various cellular processes such as cell survival, growth, proliferation and metabolism [109]. Akt1 plays a key role in inhibiting apoptosis, which is the programmed cell death that occurs when cells are damaged or no longer needed [110]. Apoptosis is regulated by a balance between pro-apoptotic and anti-apoptotic factors, and Akt1 can modulate this balance by phosphorylating and inactivating several components of the apoptotic machinery [111].

One of the targets of Akt1 is BAD, a pro-apoptotic member of the Bcl-2 family that forms heterodimers with anti-apoptotic proteins such as Bcl-XL and prevents them from blocking apoptosis [112]. When Akt1 phosphorylates BAD, it causes BAD to dissociate from Bcl-XL and bind to 14-3-3, a cytoplasmic protein that sequesters BAD from the mitochondria, where apoptosis is initiated [112]. This allows Bcl-XL to inhibit apoptosis by preventing the release of cytochrome c from the mitochondria [112].

Another target of Akt1 is IKK, a kinase that activates NF-kappaB, a transcription factor that regulates the expression of many genes involved in cell survival, inflammation and immunity [112]. Akt1 phosphorylates and activates IKK, which then phosphorylates I-kappaB, an inhibitor of NF-kappaB [112]. This leads to the degradation of I-kappaB and the release of NF-kappaB, which translocates to the nucleus and activates the transcription of anti-apoptotic genes such as Bcl-2, Bcl-XL, XIAP and c-IAP [112]. Akt1 also phosphorylates and inactivates several other pro-apoptotic factors, such as caspase-9, forkhead transcription factors and p53 [111]. By doing so, Akt1 prevents the activation of the caspase cascade, the transcription of pro-apoptotic genes and the induction of DNA damage response [111].

In summary, Akt1 is a very important kinase in the PI3K/Akt signaling pathway that promotes cell survival by inhibiting apoptosis through multiple mechanisms. Akt1 can phosphorylate and regulate various components of the apoptotic machinery, such as BAD, IKK, NF-kappaB, caspase-9, forkhead transcription factors and p53. Akt1 is essential for the normal development and function of the nervous system and other tissues [110] [113] [114].

#### 1.3. Most important Edges

1. **Septin4 (-) XIAP:** One of the most important Edges is between Septin4 and XIAP, XIAP (X-linked inhibitor of apoptosis protein) is a member of the inhibitor of apoptosis protein (IAP) family that can suppress caspase activity and regulate various cellular signaling pathways [73]. Septin4 is a protein localized at the mitochondrion that can promote cell death mainly by binding and inhibiting XIAP, thus activating caspases [74]. The interaction between XIAP and Septin4 has been implicated in several biological processes, such as apoptosis [75], cancer cell death [76], and neuronal differentiation [77]. In apoptosis, Septin4 enhances the binding between HIF-1α (hypoxia-inducible factor 1 alpha) and the E3 ubiquitin ligase VHL (von Hippel-Lindau protein) to down-regulate HIF-1α, which is a protective factor against hypoxia-induced apoptosis [75]. By reducing HIF-1α levels, Septin4 aggravates the hypoxia-induced apoptosis. Septin4 also interacts with XIAP at the Septin4-GTPase domain to promote its degradation [75].

In cancer cell death, Septin4 translocates from the mitochondria to the cytoplasm and directly binds to XIAP, leading to caspase activation and cell death [76]. Septin4 also mediates the interaction between BCL2 (B-cell lymphoma 2) and XIAP, thereby positively regulating the ubiquitination and degradation of BCL2 and promoting apoptosis [78]. BCL2 is an anti-apoptotic protein that can inhibit the release of cytochrome c from the mitochondria and prevent caspase activation [79]. In neuronal differentiation, Septin4 regulates the expression of XIAP and its downstream target genes, such as BIRC5 (baculoviral IAP repeat containing 5) and CCND1 (cyclin D1), which are involved in cell cycle progression and survival [77]. Septin4 also modulates the activity of JNK (c-Jun N-terminal kinase), a mitogen-activated protein kinase that can induce apoptosis or differentiation depending on the cellular context [80]. Septin4 promotes neuronal differentiation by inhibiting XIAP-mediated JNK activation and inducing JNK-mediated XIAP degradation [77].

#### 1.4. Enrichment analysis

1. **TNF signaling pathway**: have very important role in Apoptosis signaling pathway. The role and intersection of TNF signaling pathway in apoptosis signaling pathway is a complex and dynamic topic that has been extensively studied in various physiological and pathological contexts. TNF (tumor necrosis factor) is a pro-inflammatory cytokine that can bind to two types of receptors, TNFR1 and TNFR2, on the cell surface. Depending on the cell type, the receptor subtype, and the downstream signaling molecules, TNF can trigger different cellular responses, such as proliferation, differentiation, inflammation, and apoptosis [81]. Apoptosis is a programmed cell death process that is essential for maintaining tissue homeostasis and eliminating damaged or unwanted cells. Apoptosis can be initiated by two main pathways: the intrinsic pathway, which is mediated by mitochondrial events, and the extrinsic pathway, which is mediated by death receptors [82]. The TNF signaling pathway can interact with both pathways and modulate the balance between cell survival and death. The activation of TNFR1 by TNF leads to the recruitment of several adaptor proteins, such as TRADD, FADD, RIP, and MADD, which form a complex called TNFR1-associated death domain (TNFR1-DD). This complex can activate two distinct signaling branches: one that involves the activation of caspase-8 and leads to apoptosis, and another that involves the activation of NF-κB and leads to survival [81]. Caspase-8 is a key initiator of the extrinsic apoptosis pathway, which can directly cleave and activate downstream effector caspases, such as caspase-3, -6, and -7, or indirectly activate the intrinsic pathway by cleaving BID, a pro-apoptotic member of the Bcl-2 family. BID can translocate to the mitochondria and induce the release of cytochrome c, which forms a complex with APAF-1 and caspase-9, called apoptosome. The apoptosome can then activate caspase-3 and other effector caspases [82]. NF-κB is a transcription factor that can induce the expression of anti-apoptotic genes, such as Bcl-2, Bcl-xL, c-IAPs, FLIPs, and TRAFs, which can inhibit the activation of caspases or block the mitochondrial permeabilization [81]. The activation of TNFR2 by TNF leads to the recruitment of TRAF2 and TRAF5 proteins, which can activate several kinases, such as NIK, IKKs, MEKKs, ASK1s, JNKs, p38 MAPKs, ERKs, and AKT. These kinases can regulate various cellular processes, such as inflammation, differentiation, proliferation, and survival. Some of these kinases can also modulate apoptosis by phosphorylating pro- or anti-apoptotic proteins [81]. For example, AKT can phosphorylate and inactivate BAD and BAX, two pro-apoptotic members of the Bcl-2 family that promote mitochondrial permeabilization. AKT can also phosphorylate and activate IKKs, which can activate NF-κB [83]. JNKs can phosphorylate and activate BIM and BAX or inhibit Bcl-2 and Bcl-xL [84]. The role and intersection of TNF signaling pathway in apoptosis signaling pathway depends on the cellular context and the balance between pro- and anti-apoptotic signals. TNF can induce apoptosis in some cell types or conditions but promote survival in others. TNF can also have dual effects on the same cell type depending on the duration or intensity of its stimulation. For example, TNF can induce neuronal apoptosis in the late stage of embryonic development but increase neuronal numbers in the early stage by activating NF-κB [85]. Therefore, understanding the molecular mechanisms underlying the regulation of TNF signaling pathway in apoptosis signaling pathway is crucial for elucidating its physiological functions and pathological implications.
2. **HIV1 and hepatitis C virus (HCV):** are two major causes of chronic viral infections that can lead to serious liver diseases and increase the risk of hepatocellular carcinoma. Both viruses can interfere with the host cell apoptosis signaling pathways, which are essential for maintaining the balance between cell survival and death. Apoptosis can be triggered by two main pathways: the extrinsic pathway, which involves the activation of death receptors such as Fas or TNF-α on the cell surface, and the intrinsic pathway, which involves the release of cytochrome c from the mitochondria and the activation of caspases. Both pathways converge on the activation of caspase-3, which executes the apoptotic program by cleaving various cellular substrates [86].

HIV and HCV can modulate the apoptosis signaling pathways in different ways, depending on the stage of infection, the cell type, and the viral genotype. Some studies have shown that HIV and HCV can induce apoptosis in infected cells or bystander cells by upregulating the expression of pro-apoptotic molecules such as FasL, TNF-α, or Bcl-2 family members [87][88][89]. This may help the viruses to evade the immune system, to spread to new cells, or to create a pro-inflammatory environment that favors viral replication. On the other hand, some studies have shown that HIV and HCV can inhibit apoptosis in infected cells by downregulating the expression of pro-apoptotic molecules or by activating anti-apoptotic pathways such as NF-κB, PI3K/Akt, or STAT3 [87][88][89]. This may help the viruses to establish persistent infection, to avoid clearance by cytotoxic T cells, or to promote cell survival and proliferation. One of the key regulators of apoptosis signaling pathways in HIV/HCV coinfected cells is signal transducer and activator of transcription factor 1 (STAT1), which is activated by interferons and other cytokines. STAT1 can induce the expression of various interferon-stimulated genes (ISGs) that have antiviral and pro-apoptotic effects, such as CXCL10, MX1, IFIT1, or IRG1 [90]. However, HIV and HCV can also interfere with STAT1 signaling by blocking its phosphorylation, nuclear translocation, or DNA binding [90]. Moreover, HIV and HCV can modulate the expression of other STAT family members, such as STAT3 or STAT5, which have anti-apoptotic and pro-survival effects [90]. The intersection of HIV and HCV signaling pathways with apoptosis signaling pathway is a complex and dynamic process that involves multiple factors and feedback loops.

### 2. Cell-cycle signaling pathway Algorithm

(Precision: 97%).

Similar to (Figure 3), every node in (Figure 7) has interaction with other nodes according to the weight of edges. If the relationship between two nodes is higher, then they will have a higher edge weight. In our Hierarchic layout, if one edge has a higher weight, then that edge has higher thickness and higher color contrast in our Hierarchic layout.

**Figure 6.**
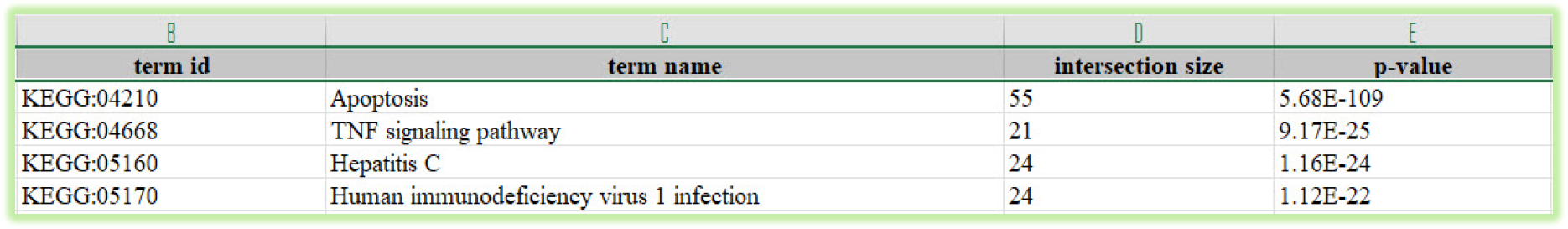
Enrichment analysis of Apoptosis signaling pathway.

**Figure 7.**
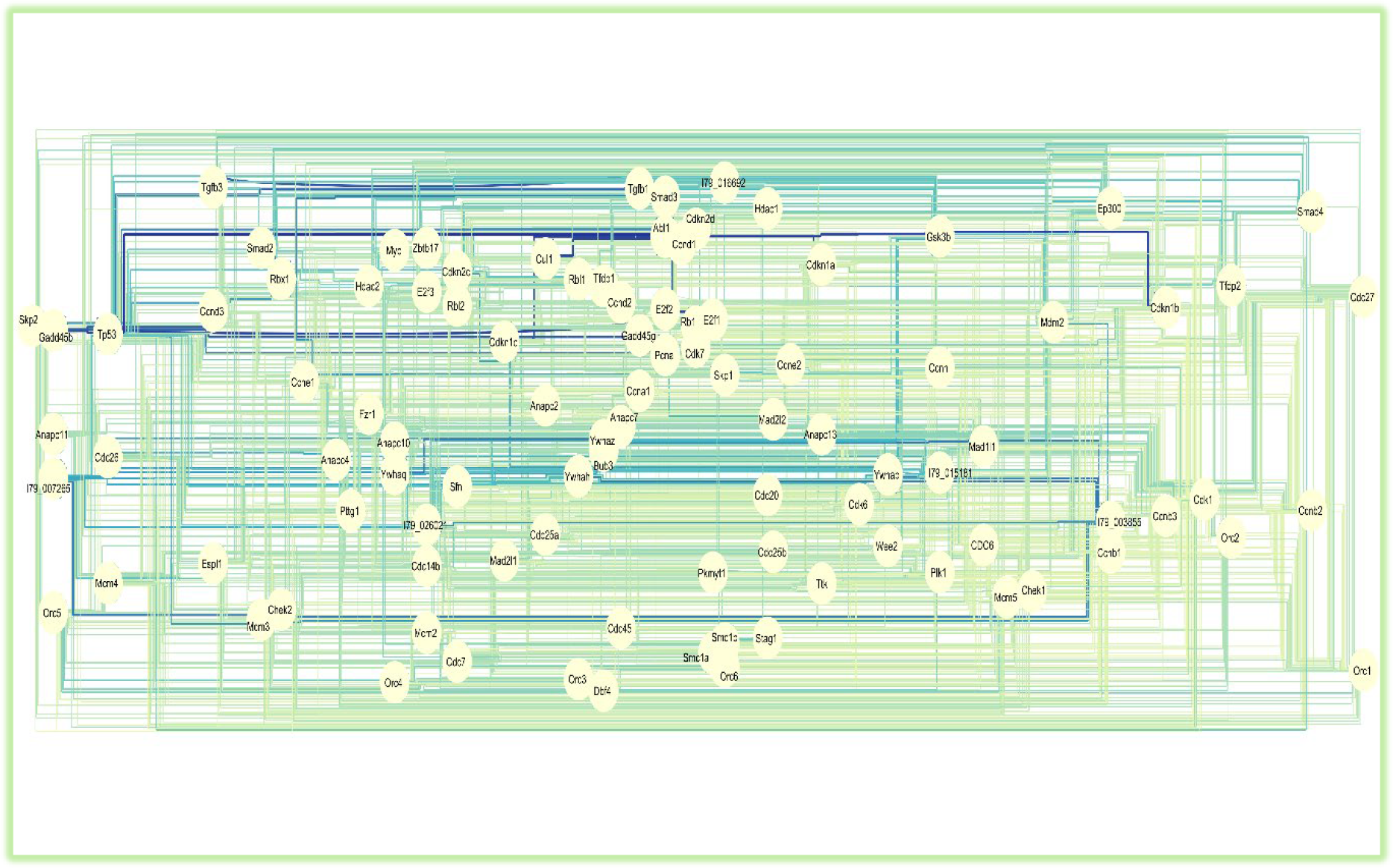
Hierarchic Layout of GAT Cell-cycle signaling pathway Algorithm.

**Figure 8.**
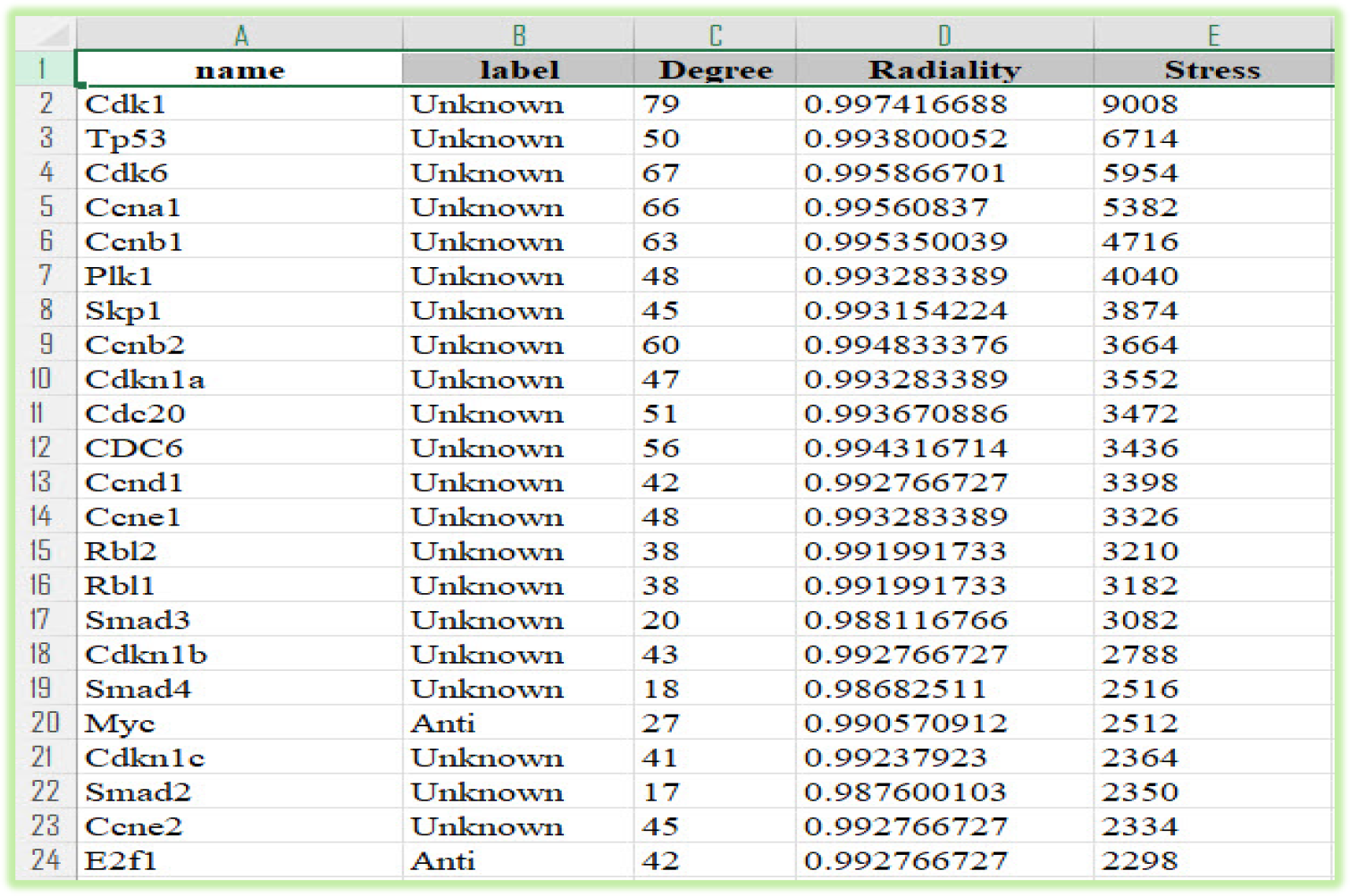
Most important Nodes (proteins) in Cell-cycle signaling pathway.

**Figure 9.**
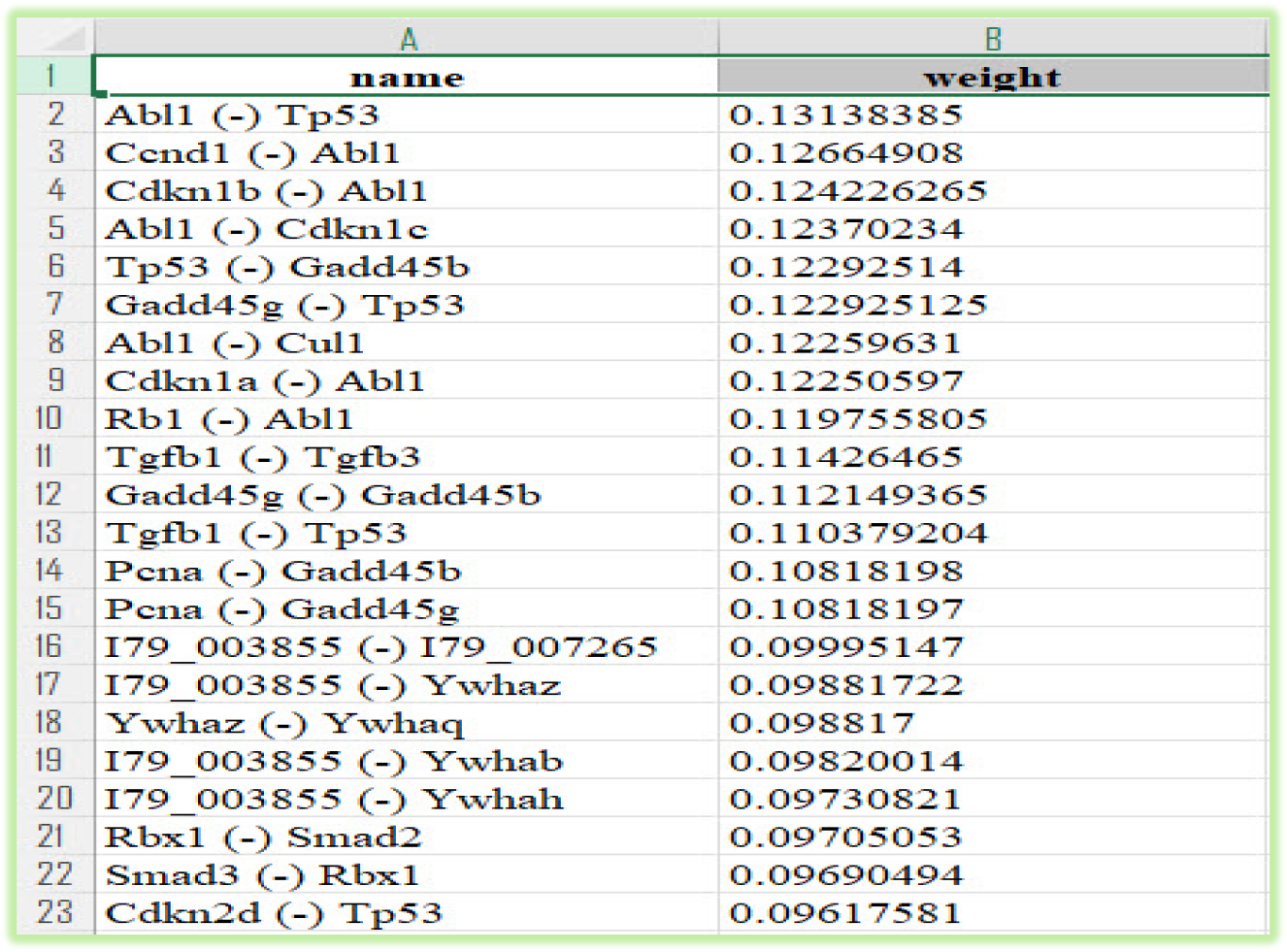
Most important Edges (Interactions between proteins) in Cell-cycle signaling pathway.

#### 2.1. Most important Nodes

1. **Cdk1:** One of the most important Nodes (proteins) in Cell-cycle signaling pathway that have interaction with Apoptosis pathway is Cdk1. Cdk1 is a cyclin-dependent kinase that plays a key role in regulating the cell cycle and the interaction of Cdk1 with apoptosis pathway. Cdk1 stimulates the progression from G2 to M phase of the cell cycle by phosphorylating various substrates, such as nuclear lamins, histone H1, and microtubule-associated proteins [91]. Cdk1 also interacts with the apoptosis pathway in different ways, depending on the cellular context and the level of stress or DNA damage [92].

One of the mechanisms by which Cdk1 can induce apoptosis is by blocking the activity of TP53, a tumor suppressor protein that can trigger cell cycle arrest or apoptosis in response to DNA damage [93]. Cdk1 can phosphorylate TP53 at Ser315, which prevents its translocation to the nucleus and its transcriptional activation of pro-apoptotic genes [94]. Alternatively, Cdk1 can activate E2F1, a transcription factor that can induce both cell cycle progression and apoptosis [95]. E2F1 can upregulate the expression of pro-apoptotic genes, such as BAX, BID, and PUMA, and also downregulate the expression of anti-apoptotic genes, such as BCL-2 and BCL-XL [96]. Another mechanism by which Cdk1 can modulate apoptosis is by interacting with the PANoptosome, a multiprotein complex that mediates pyroptosis, apoptosis, and necroptosis (PANoptosis) [97]. Cdk1 can bind to the PANoptosome components, such as CASP1, CASP8, RIPK3, and ZBP1, and regulate their activation and cleavage [97]. Cdk1 can also phosphorylate ZBP1 at Ser181, which enhances its binding to RIPK3 and promotes necroptosis [97]. Moreover, Cdk1 can inhibit pyroptosis by phosphorylating NLRP3 at Ser5, which impairs its oligomerization and inflammasome formation [97].

In summary, Cdk1 is a critical regulator of the cell cycle and the interaction of Cdk1 with apoptosis pathway. Cdk1 can modulate the balance between cell survival and death by influencing different signaling pathways and molecular targets. The dysregulation of Cdk1 activity or expression can contribute to tumorigenesis and cancer progression by enhancing cell proliferation and evading apoptosis.

3. **Cdk6:** Cdk6 is a cyclin-dependent kinase that regulates the cell cycle progression from G1 to S phase. It forms a complex with cyclin D and phosphorylates the retinoblastoma protein (Rb), which releases the transcription factor E2F to activate genes required for DNA synthesis [115]. Cdk6 also has other roles in cell fate determination, such as regulating stem cell quiescence, nuclear factor kappaB (NF-κB) signaling, and apoptosis [116].

Cdk6 is involved in the cell-cycle signaling pathway that controls the balance between cell proliferation and cell death. When Cdk6 is overexpressed or deregulated, it can lead to abnormal cell growth and cancer development [117]. Therefore, Cdk6 is a potential target for cancer therapy, especially for tumors that are resistant to conventional treatments. For example, Cdk6 inhibition can attenuate the expression and function of ABCB1/P-glycoprotein, a multidrug resistance (MDR) transporter that pumps out chemotherapeutic agents from cancer cells [118]. Cdk6 inhibition can also induce alternative splicing (AS) of ABCB1 pre-mRNA, which results in lower levels of functional ABCB1 protein [118].

Cdk6 also interacts with the apoptosis signaling pathway, which is responsible for triggering programmed cell death in response to various stimuli, such as DNA damage, oxidative stress, or cytotoxic drugs. Cdk6 can modulate the accumulation of pro-apoptotic proteins, such as p53 and p130, which prevent cells from entering the cell cycle if there is DNA damage and activate the intrinsic or extrinsic apoptotic pathways [119]. Cdk6 inhibition can enhance the sensitivity of cancer cells to apoptosis by downregulating anti-apoptotic factors, such as Bcl-2 and survivin, and upregulating pro-apoptotic factors, such as Bax and caspase-3 [120].

In conclusion, Cdk6 is a key regulator of the cell-cycle signaling pathway and its relation with the apoptosis signaling pathway. It plays an important role in determining the fate of cells, either proliferation or death. Cdk6 is also a promising target for cancer therapy, as it can overcome MDR and induce apoptosis in cancer cells.

#### 2.2. Most important Edges

1. **Abl1 (-) Tp53**: One the most important Edges between Abl1 and Tp53. The interaction between Abl1 and Tp53 is an important regulator of cell cycle and apoptosis signaling pathways. Abl1 is a tyrosine kinase that can phosphorylate Tp53 on several residues, modulating its activity and stability. Abl1 can activate the proapoptotic pathway when the DNA damage is too severe to be repaired, by phosphorylating Tp73, a homolog of Tp53, and caspase-9, an initiator of the intrinsic apoptotic pathway. Abl1 can also inhibit the antiapoptotic pathway by phosphorylating Mdm2, an E3 ubiquitin ligase that targets Tp53 for degradation [98]. Tp53 is a transcription factor that responds to various cellular stresses, such as DNA damage, hypoxia, and oncogene activation, by activating genes involved in cell cycle arrest, DNA repair, senescence, and apoptosis. Tp53 can trigger the extrinsic apoptotic pathway by inducing the expression of Fas and DR5, two death receptors that bind to FasL and TRAIL, respectively, and activate caspase-8. Tp53 can also trigger the intrinsic apoptotic pathway by inducing the expression of Bax, Puma, Noxa, and Apaf-1, which promote the release of cytochrome c from the mitochondria and the formation of the apoptosome, a complex that activates caspase-9 [99,100]. The interaction between Abl1 and Tp53 is crucial for maintaining genomic integrity and preventing tumorigenesis. Mutations or dysregulation of either gene can impair the apoptotic response to DNA damage and lead to cancer development. For example, Abl1 is involved in the chromosomal translocation that causes chronic myeloid leukemia (CML), resulting in a constitutively active fusion protein with Bcr that promotes cell proliferation and survival. Tp53 is one of the most frequently mutated genes in human cancers, and its loss or inactivation confers resistance to chemotherapy and radiation therapy [101,102,103].
2. **Ccnd1 (-) Abl1:** The interaction between Ccnd1 and Abl1 is a key factor in the regulation of cell cycle and apoptosis signaling pathways. Ccnd1, also known as cyclin D1, is a proto-oncogene that forms active complexes with CDK4 and CDK6, promoting the transition from G1 to S phase by phosphorylating and inactivating the retinoblastoma protein (RB) [121]. Abl1, also known as c-Abl, is a non-receptor tyrosine kinase that has multiple roles in cellular processes, including DNA damage response, cytoskeleton remodeling, cell adhesion and apoptosis [122].

The balance between Ccnd1 and Abl1 activities determines the fate of the cell in response to various stimuli. Under normal conditions, Ccnd1 expression is tightly regulated by transcriptional and post-translational mechanisms, ensuring proper cell cycle progression [123]. However, in many cancers, Ccnd1 is overexpressed or amplified, leading to uncontrolled proliferation and resistance to apoptosis [124]. Abl1 can counteract the oncogenic effects of Ccnd1 by inducing its degradation via the ubiquitin pathway [125]. This process requires an RxxL destruction box within the N-terminus of Ccnd1 and is triggered by genotoxic stress, such as ionizing radiation or DNA-damaging agents [125]. Abl1 can also induce apoptosis by activating the intrinsic mitochondrial pathway or the extrinsic death receptor pathway, depending on the cellular context [122].

Therefore, the interaction between Ccnd1 and Abl1 is crucial for maintaining cellular homeostasis and preventing tumorigenesis. Targeting this interaction with specific inhibitors or modulators may offer novel therapeutic strategies for cancer treatment. For example, combining Abl inhibitors with Src inhibitors has been shown to enhance apoptosis, alter cell cycle regulation and reduce CD34+ cells in BCR-ABL1 positive leukemia cells [126]. Moreover, manipulating the expression or activity of Ccnd1 may affect cellular metabolism and sensitize cancer cells to fatty acid oxidation inhibitors [127].

## Conclusion and Discussion

In this paper, we introduce the GAT machine learning algorithm that can focus on and find the most important targets in the process of CHO cell death. In previous works, only two or three targets were analyzed, and the interaction of these proteins was ignored. To analyze these complex signaling pathways, we need to examine all components of the signaling pathways and identify the most important targets. Advanced machine learning algorithms are the best way to find these targets. Traditional bioinformatics tools cannot analyze these large and complex pathways, so we require modern and high-performance methods.

In our analysis, Tp53 is identified as the most important target in both the apoptosis and cell-cycle signaling pathways, with a high weight of edges connecting it to all other nodes in the pathways. In the cell signaling pathways section, the TNF signaling pathway has the most intersection with the apoptosis signaling pathway. Additionally, many nodes in the TNF signaling pathway play a role in the apoptosis pathway. Some of these targets have been validated with experiments in the laboratory in previous works, but they have not been compared to each other to determine which one is more efficient. Therefore, advanced laboratories are needed to validate these targets and compare them. High-throughput screening is one of the most important methods that we can use to experiment with these targets. Advanced AI solutions, like the one presented in this work, can significantly accelerate the experimental validation of these targets.

## References

[1] K. P. Jayapal, K. F. Wlaschin, W. S. Hu, and M. J. Yap, “Recombinant protein therapeutics from CHO cells : 20 years and counting,” Chemical Engineering Progress, vol. 103, no. 10, pp. 40–47, Oct. 2007, [Online]. Available: https://experts.umn.edu/en/publications/recombinant-protein-therapeutics-from-cho-cells-20-years-and-coun.

[2] M. Pereira, “CHO in biomanufacturing: Past, present and future,” European Pharmaceutical Manufacturer, 01-Jun-2021. [Online]. Available: https://pharmaceuticalmanufacturer.media/pharmaceutical-industry-insights/biopharma-news/cho-in-biomanufacturing-past-present-and-future. [Accessed: 02-Dec-2023].

[3] “HEK cells vs. CHO cells in recombinant antibody production: What’s the better choice,” Pharmiweb.com, 03-Mar-2022. [Online]. Available: https://www.pharmiweb.com/article/hek-cells-vs-cho-cells-in-recombinant-antibody-production-whats-the-better-choice. [Accessed: 02-Dec-2023].

[4] Keysberg, C., Hertel, O., Schelletter, L., Busche, T., Sochart, C., Kalinowski, J., … & Noll, T. (2021). Exploring the molecular content of CHO exosomes during bioprocessing. Applied microbiology and biotechnology, 105, 3673–3689.

[5] Xu, X., Nagarajan, H., Lewis, N. E., Pan, S., Cai, Z., Liu, X., … & Wang, J. (2011). The genomic sequence of the Chinese hamster ovary (CHO)-K1 cell line. Nature biotechnology, 29(8), 735–741.

[6] Kildegaard, H. F., Baycin-Hizal, D., Lewis, N. E., & Betenbaugh, M. J. (2013). The emerging CHO systems biology era: harnessing the ‘omics revolution for biotechnology. Current opinion in biotechnology, 24(6), 1102–1107.

[7] Lee, J. S., Grav, L. M., Lewis, N. E., & Faustrup Kildegaard, H. (2015). CRISPR/Cas9-mediated genome engineering of CHO cell factories: Application and perspectives. Biotechnology journal, 10(7), 979–994.

[8] GlobalData UK Ltd., “Top 20 global biopharmaceutical companies report marginal growth in market cap during Q3 2023, reveals GlobalData,” GlobalData UK Ltd, 30-Oct-2023. [Online]. Available: https://www.globaldata.com/media/business-fundamentals/top-20-global-biopharmaceutical-companies-report-marginal-growth-in-market-cap-during-q3-2023-reveals-globaldata/. [Accessed: 02-Dec-2023].

[9] Wurm, F. M. (2004). Production of recombinant protein therapeutics in cultivated mammalian cells. Nature biotechnology, 22(11), 1393–1398.

[10] Kim, J. Y., Kim, Y. G., & Lee, G. M. (2012). CHO cells in biotechnology for production of recombinant proteins: current state and further potential. Applied microbiology and biotechnology, 93, 917–930.

[11] Fussenegger, M., & Bailey, J. E. (1998). Molecular regulation of cell-cycle progression and apoptosis in mammalian cells: implications for biotechnology. Biotechnology progress, 14(6), 807–833.

[12] Arnold, F. H. (1998). Design by directed evolution. Accounts of chemical research, 31(3), 125–131.

[13] Walsh, G. (2018). Biopharmaceutical benchmarks 2018. Nature biotechnology, 36(12), 1136–1145.

[14] Majors, B. S., Betenbaugh, M. J., & Chiang, G. G. (2007). Links between metabolism and apoptosis in mammalian cells: applications for anti-apoptosis engineering. Metabolic engineering, 9(4), 317–326.

[15] Elmore, S. (2007). Apoptosis: a review of programmed cell death. Toxicologic pathology, 35(4), 495–516.

[16] Peter, M. E., & Krammer, P. H. (2003). The CD95 (APO-1/Fas) DISC and beyond. Cell Death & Differentiation, 10(1), 26–35.

[17] Lavrik, I. N., Golks, A., & Krammer, P. H. (2005). Caspases: pharmacological manipulation of cell death. The Journal of clinical investigation, 115(10), 2665–2672.

[18] Green, D. R., & Kroemer, G. (2004). The pathophysiology of mitochondrial cell death. Science, 305(5684), 626–629.

[19] Li, P., Nijhawan, D., Budihardjo, I., Srinivasula, S. M., Ahmad, M., Alnemri, E. S., & Wang, X. (1997). Cytochrome c and dATP-dependent formation of Apaf-1/caspase-9 complex initiates an apoptotic protease cascade. cell, 91(4), 479–489.

[20] Pop, C., & Salvesen, G. S. (2009). Human caspases: activation, specificity, and regulation. Journal of biological Chemistry, 284(33), 21777–21781.

[21] Grilo, A. L., & Mantalaris, A. (2019). Apoptosis: A mammalian cell bioprocessing perspective. Biotechnology advances, 37(3), 459–475.

[22] Evan, G. I., & Vousden, K. H. (2001). Proliferation, cell cycle and apoptosis in cancer. nature, 411(6835), 342–348.

[23] Bryan, L., Henry, M., Kelly, R.M. et al. Mapping the molecular basis for growth related phenotypes in industrial producer CHO cell lines using differential proteomic analysis. BMC Biotechnol 21, 43 (2021). 10.1186/s12896-021-00704-8

[24] Lewis, N. E., Liu, X., Li, Y., Nagarajan, H., Yerganian, G., O’brien, E., … & Palsson, B. O. (2013). Genomic landscapes of Chinese hamster ovary cell lines as revealed by the Cricetulus griseus draft genome. Nature biotechnology, 31(8), 759–765.

[25] Yang, F., Fan, K., Song, D., & Lin, H. (2020). Graph-based prediction of protein-protein interactions with attributed signed graph embedding. BMC bioinformatics, 21(1), 1–16.

[26] Lai, T., Yang, Y., & Ng, S. K. (2013). Advances in mammalian cell line development technologies for recombinant protein production. Pharmaceuticals, 6(5), 579–603.

[27] Lv, G., Hu, Z., Bi, Y., & Zhang, S. (2021). Learning unknown from correlations: graph neural network for inter-novel-protein interaction prediction. arXiv preprint arXiv:2105.06709.

[28] Wu, Z., Pan, S., Chen, F., Long, G., Zhang, C., & Philip, S. Y. (2020). A comprehensive survey on graph neural networks. IEEE transactions on neural networks and learning systems, 32(1), 4–24.

[29] Velickovic, P., Cucurull, G., Casanova, A., Romero, A., Lio, P., & Bengio, Y. (2017). Graph attention networks. arXiv preprint arXiv:1710.10903.

[30] Data Camp (2021) A Comprehensive Introduction to Graph Neural Networks (GNNs). https://www.datacamp.com/tutorial/comprehensive-introduction-graph-neural-networks-gnns-tutorial.

[31] “pandas,” Pydata.org. [Online]. Available: https://pandas.pydata.org/. [Accessed: 02-Dec-2023].

[32] “NumPy,” Numpy.org. [Online]. Available: https://numpy.org/. [Accessed: 02-Dec-2023].

[33] Pytorch.org. [Online]. Available: https://pytorch.org/. [Accessed: 02-Dec-2023].

[34] “PyG Documentation — pytorch_geometric documentation,” Readthedocs.io. [Online]. Available: https://pytorch-geometric.readthedocs.io/en/latest/. [Accessed: 02-Dec-2023].

[35] “Scikit-learn,” Scikit-learn.org. [Online]. Available: https://scikit-learn.org/stable/. [Accessed: 02-Dec-2023].

[36] “NetworkX — NetworkX documentation,” Networkx.org. [Online]. Available: https://networkx.org/. [Accessed: 02-Dec-2023].

[37] “JetBrains DataSpell: The IDE for data scientists,” JetBrains. [Online]. Available: https://www.jetbrains.com/dataspell/. [Accessed: 02-Dec-2023].

[38] K. Ono, “Cytoscape,” Cytoscape.org. [Online]. Available: https://cytoscape.org/. [Accessed: 02-Dec-2023].

[39] “Sublime Text - the sophisticated text editor for code, markup and prose,” Sublimetext.com. [Online]. Available: https://www.sublimetext.com/. [Accessed: 02-Dec-2023].

[40] Anaconda.com. [Online]. Available: https://www.anaconda.com/. [Accessed: 02-Dec-2023].

[41] “STRING: functional protein association networks,” String-db.org. [Online]. Available: https://string-db.org/. [Accessed: 02-Dec-2023].

[42] “KEGG: Kyoto Encyclopedia of Genes and genomes,” Genome.jp. [Online]. Available:https://www.genome.jp/kegg/. [Accessed: 02-Dec-2023].

[43] MacDonald, M. A., Barry, C., Groves, T., Martínez, V. S., Gray, P. P., Baker, K., … & Nielsen, L. K. (2022). Modeling apoptosis resistance in CHO cells with CRISPR-mediated knockouts of Bak1,Bax, and Bok. Biotechnology and Bioengineering, 119(6), 1380–1391.

[44] Cost, G. J., Freyvert, Y., Vafiadis, A., Santiago, Y., Miller, J. C., Rebar, E., … & Gregory, P. D. (2010). BAK and BAX deletion using zinc-finger nucleases yields apoptosis-resistant CHO cells. Biotechnology and bioengineering, 105(2), 330–340.

[45] Henry, M. N., MacDonald, M. A., Orellana, C. A., Gray, P. P., Gillard, M., Baker, K., … & Martínez, V. S. (2020). Attenuating apoptosis in Chinese hamster ovary cells for improved biopharmaceutical production. Biotechnology and Bioengineering, 117(4), 1187–1203.

[46] Orellana, C. A., Martínez, V. S., MacDonald, M. A., Henry, M. N., Gillard, M., Gray, P. P., … & Marcellin, E. (2021). ‘Omics driven discoveries of gene targets for apoptosis attenuation in CHO cells. Biotechnology and Bioengineering, 118(1), 481–490.

[47] Lee, J. S., Ha, T. K., Park, J. H., & Lee, G. M. (2013). Anti-cell death engineering of CHO cells: Co-overexpression of Bcl-2 for apoptosis inhibition, Beclin-1 for autophagy induction. Biotechnology and bioengineering, 110(8), 2195–2207.

[48] Krampe, B., & Al-Rubeai, M. (2010). Cell death in mammalian cell culture: molecular mechanisms and cell line engineering strategies. Cytotechnology, 62, 175–188.

[49] Arden, N., & Betenbaugh, M. J. (2004). Life and death in mammalian cell culture: strategies for apoptosis inhibition. TRENDS in Biotechnology, 22(4), 174–180.

[50] Arden, N., & Betenbaugh, M. J. (2006). Regulating apoptosis in mammalian cell cultures. Cytotechnology, 50, 77–92.

[51] Tang, D., Lam, C., Bauer, N., Auslaender, S., Snedecor, B., Laird, M. W., & Misaghi, S. (2022). Bax and Bak knockout apoptosis-resistant Chinese hamster ovary cell lines significantly improve culture viability and titer in intensified fed-batch culture process. Biotechnology Progress, 38(2), e3228.

[52] Figueroa Jr, B., Sauerwald, T. M., Oyler, G. A., Hardwick, J. M., & Betenbaugh, M. J. (2003). A comparison of the properties of a Bcl-xL variant to the wild-type anti-apoptosis inhibitor in mammalian cell cultures. Metabolic Engineering, 5(4), 230–245.

[53] Figueroa Jr, B., Chen, S., Oyler, G. A., Hardwick, J. M., & Betenbaugh, M. J. (2004). Aven and Bcl-xL enhance protection against apoptosis for mammalian cells exposed to various culture conditions. Biotechnology and bioengineering, 85(6), 589–600.

[54] Sauerwald, T. M., Figueroa Jr, B., Hardwick, J. M., Oyler, G. A., & Betenbaugh, M. J. (2006). Combining caspase and mitochondrial dysfunction inhibitors of apoptosis to limit cell death in mammalian cell cultures. Biotechnology and bioengineering, 94(2), 362–372.

[55] Albrecht, S., Kaisermayer, C., Gallagher, C., Farrell, A., Lindeberg, A., & Bones, J. (2018). Proteomics in biomanufacturing control: Protein dynamics of CHO-K1 cells and conditioned media during apoptosis and necrosis. Biotechnology and Bioengineering, 115(6), 1509–1520.

[56] Arden, N., Majors, B. S., Ahn, S. H., Oyler, G., & Betenbaugh, M. J. (2007). Inhibiting the apoptosis pathway using MDM2 in mammalian cell cultures. Biotechnology and bioengineering, 97(3), 601–614.

[57] Zhao, X., Guo, J., Yu, Y., Yi, S., Yu, T., Fu, L., … & Chen, W. (2011). Overexpression of survivin and cyclin D1 in CHO cells confers apoptosis resistance and enhances growth in serum-free suspension culture. Biotechnology letters, 33, 1293–1300.

[58] Liew, J. C., Tan, W. S., Alitheen, N. B. M., Chan, E. S., & Tey, B. T. (2010). Over-expression of the X-linked inhibitor of apoptosis protein (XIAP) delays serum deprivation-induced apoptosis in CHO-K1 cells. Journal of bioscience and bioengineering, 110(3), 338–344.

[59] Skretting, G., Iversen, N., Myklebust, C. F., Dahm, A. E., & Sandset, P. M. (2012). Overexpression of tissue factor pathway inhibitor in CHO-K1 cells results in increased activation of NF-κB and apoptosis mediated by a caspase-3 independent pathway. Molecular biology reports, 39, 10089–10096.

[60] Fan, W., Ha, T., Li, Y., Ozment-Skelton, T., Williams, D. L., Kelley, J., … & Li, C. (2005). Overexpression of TLR2 and TLR4 susceptibility to serum deprivation-induced apoptosis in CHO cells. Biochemical and biophysical research communications, 337(3), 840–848.

[61] Ifandi, V., & Al-Rubeai, M. (2005). Regulation of cell proliferation and apoptosis in CHO-K1 cells by the coexpression of c-Myc and Bcl-2. Biotechnology progress, 21(3), 671–677.

[62] Wong, D. C. F., Wong, K. T. K., Lee, Y. Y., Morin, P. N., Heng, C. K., & Yap, M. G. S. (2006). Transcriptional profiling of apoptotic pathways in batch and fed-batch CHO cell cultures. Biotechnology and bioengineering, 94(2), 373–382.

[63] Rupp, O., MacDonald, M. L., Li, S., Dhiman, H., Polson, S., Griep, S., … & Lee, K. H. (2018). A reference genome of the Chinese hamster based on a hybrid assembly strategy. Biotechnology and bioengineering, 115(8), 2087–2100.

[64] Baycin-Hizal, D., Tabb, D. L., Chaerkady, R., Chen, L., Lewis, N. E., Nagarajan, H., … & Betenbaugh, M. (2012). Proteomic analysis of Chinese hamster ovary cells. Journal of proteome research, 11(11), 5265–5276.

[65] Baeldung.com. [Online]. Available: https://www.baeldung.com/cs/cross-validation-k-fold-loo. [Accessed: 02-Dec-2023].

[66] Zach, “A quick intro to leave-one-out cross-validation (LOOCV),” Statology, 03-Nov-2020. [Online]. Available: https://www.statology.org/leave-one-out-cross-validation/. [Accessed: 02-Dec-2023].

[67] N. Khadka, “How Leave-one-out Cross Validation (LOOCV) improve’s model performance,” Dataaspirant - A Data Science Portal For Beginners, 05-Oct-2023.

[68] “g:Profiler – a web server for functional enrichment analysis and conversions of genelists,” Cs.ut.ee. [Online]. Available: https://biit.cs.ut.ee/gprofiler/page/apis. [Accessed: 02-Dec-2023].

[69] Reimand J, Kull M, Peterson H, Hansen J, Vilo J. g:Profiler--a web-based toolset for functional profiling of gene lists from large-scale experiments. Nucleic Acids Res. 2007 Jul;35(Web Server issue):W193–200.

[70] yWorks, the diagramming experts, “yWorks - the diagramming experts,” yWorks, the diagramming experts. [Online]. Available: https://www.yworks.com/. [Accessed: 02-Dec-2023].

[71] “P53 pathway for apoptosis signaling - UK.” https://www.thermofisher.com/us/en/home/life-science/antibodies/antibodies-learning-center/antibodies-resource-library/cell-signaling-pathways/p53-mediated-apoptosis-pathway.html

[72] Marei, H. E., Althani, A., Afifi, N., Hasan, A., Caceci, T., Pozzoli, G., … & Cenciarelli, C. (2021). p53 signaling in cancer progression and therapy. Cancer cell international, 21(1), 1–15.

[73] Hanifeh, M., & Ataei, F. (2022). XIAP as a multifaceted molecule in Cellular Signaling. Apoptosis, 27(7-8), 441–453.

[74] UniProt. SEPT4_HUMAN. https://www.uniprot.org/uniprot/O43236

[75] Wu, S., Zhang, Y., You, S., Lu, S., Zhang, N., & Sun, Y. (2021). Septin4 promotes cardiomyocytes apoptosis by enhancing the VHL-mediated degradation of HIF-1α. Cell Death Discovery, 7(1), 172.

[76] Zhao, X., Feng, H., Wang, Y., Wu, Y., Guo, Q., Feng, Y., … & Cao, L. (2020). Septin4 promotes cell death in human colon cancer cells by interacting with BAX. International Journal of Biological Sciences, 16(11), 1917.

[77] Lee JH, Kim KT. Septin 4 regulates neuronal differentiation via interaction with X-linked inhibitor of apoptosis protein during mouse brain development. Cell Death Differ. 2019;26:1130–1146. https://www.nature.com/articles/s41418-018-0209-x.

[78] Liu Y, Wang Z, Wang J, Lam W, Kwong S, Li F et al. Inhibition of ubiquitin-specific protease 14 induces apoptosis through Bcl-2-mediated cross-talk between autophagy and apoptosis in human pancreatic cancer cells. Oncotarget. 2016;7:36293–36307. https://www.oncotarget.com/article/10029/text/.

[79] Youle, R. J., & Strasser, A. (2008). The BCL-2 protein family: opposing activities that mediate cell death. Nature reviews Molecular cell biology, 9(1), 47–59.

[80] Coffey, E. T. (2014). Nuclear and cytosolic JNK signaling in neurons. Nature Reviews Neuroscience, 15(5), 285–299.

[81] Thermofisher.com. [Online]. Available: https://www.thermofisher.com/us/en/home/life-science/antibodies/antibodies-learning-center/antibodies-resource-library/cell-signaling-pathways/tnf-signaling-pathway. [Accessed: 02-Dec-2023].

[82] Thermofisher.com. [Online]. Available: https://www.thermofisher.com/us/en/home/life-science/antibodies/antibodies-learning-center/antibodies-resource-library/cell-signaling-pathways/apoptosis-pathways. [Accessed: 02-Dec-2023].

[83] P. C. Rath and B. B. Aggarwal, J. Clin. Immunol., vol. 19, no. 6, pp. 350–364, 1999.

[84] K. You, H. Gu, Z. Yuan, and X. Xu, “Tumor necrosis factor alpha signaling and organogenesis,” Front. Cell Dev. Biol., vol. 9, 2021.

[85] L. L. Sandell et al., “RDH10 is essential for synthesis of embryonic retinoic acid and is required for limb, craniofacial, and organ development,” Genes Dev., vol. 21, no. 9, pp. 1113–1124, 2007.

[86] Elmore S. poptosis: a review of programmed cell death. Toxicol Pathol. 2007;35(4):495–516.

[87] Brochado-Kith O et al. HCV Cure With Direct-Acting Antivirals Improves Liver and Immunological Markers in HIV/HCV-Coinfected Patients. Front Immunol. 2021;12:723196.

[88] Li S et al. Chronic hepatitis C virus infection triggers spontaneous differential expression of biosignatures associated with T cell exhaustion and apoptosis signaling in peripheral blood mononucleocytes. Apoptosis. 2015;20(2):221–35.

[89] Khan S et al. Viral Persistence and Chronicity in Hepatitis C Virus Infection: Role of T-Cell Apoptosis, Senescence and Exhaustion. Cells. 2018;7(10):165.

[90] Ray RB et al. Signal Transducer and Activator of Transcription Factor 1 Mediates Apoptosis Induced by Hepatitis C Virus and HIV Envelope Proteins in Hepatocytes. J Infect Dis. 2006;194(5):670–81.

[91] R. Wijnen et al., “Cyclin dependent kinase-1 (CDK-1) inhibition as a novel therapeutic strategy against pancreatic ductal adenocarcinoma (PDAC),” Cancers (Basel), vol. 13, no. 17, p. 4389, 2021.

[92] L. Ren et al., “CDK1 serves as a therapeutic target of adrenocortical carcinoma via regulating epithelial–mesenchymal transition, G2/M phase transition, and PANoptosis,” J. Transl. Med., vol. 20, no. 1, 2022.

[93] R. Zhang et al., “The aberrant upstream pathway regulations of CDK1 protein were implicated in the proliferation and apoptosis of ovarian cancer cells,” J. Ovarian Res., vol. 10, no. 1, 2017.

[94] Sciencedirect.com. [Online]. Available: https://www.sciencedirect.com/science/article/pii/S0304419X16301638. [Accessed: 02-Dec-2023].

[95] Nature.com. [Online]. Available: https://www.nature.com/articles/s41419-019-1407-0. [Accessed: 02-Dec-2023].

[96] Sciencedirect.com. [Online]. Available: https://www.sciencedirect.com/science/article/pii/S016748891500053X. [Accessed: 02-Dec-2023].

[97] Nature.com. [Online]. Available: https://www.nature.com/articles/s41467-020-18276-w. [Accessed: 02-Dec-2023].

[98] “Account - GeneCards Suite,” Genecards.org. [Online]. Available: https://www.genecards.org/cgi-bin/carddisp.pl?gene=ABL1. [Accessed: 02-Dec-2023].

[99] Biologists.com. [Online]. Available: https://journals.biologists.com/jcs/article/116/20/4077/27420/Apoptosis-the-p53-network. [Accessed: 02-Dec-2023].

[100] https://www.thermofisher.com/us/en/home/life-science/antibodies/antibodies-learning-center/antibodies-resource-library/cell-signaling-pathways/apoptosis-pathways.

[101] H. E. Marei et al., “P53 signaling in cancer progression and therapy,” Cancer Cell Int., vol. 21, no. 1, 2021.

[102] https://www.thermofisher.com/us/en/home/life-science/antibodies/antibodies-learning-center/antibodies-resource-library/cell-signaling-pathways/p53-mediated-apoptosis-pathway.

[103] Mdpi.com. [Online]. Available: https://www.mdpi.com/2072-6694/10/4/110. [Accessed: 02-Dec-2023].

[104] Wikipedia contributors, “Bcl-2,” Wikipedia, The Free Encyclopedia, 19-Aug-2023. [Online]. Available: https://en.wikipedia.org/w/index.php?title=Bcl-2&oldid=1171113478.

[105] M. S. Ola, M. Nawaz, and H. Ahsan, “Role of Bcl-2 family proteins and caspases in the regulation of apoptosis,” Mol. Cell. Biochem., vol. 351, no. 1–2, pp. 41–58, 2011.

[106] F. Tzifi, C. Economopoulou, D. Gourgiotis, A. Ardavanis, S. Papageorgiou, and A. Scorilas, “The role of BCL2 family of apoptosis regulator proteins in acute and chronic leukemias,” Adv. Hematol., vol. 2012, pp. 1–15, 2012.

[107] A. W. Roberts, “Therapeutic development and current uses of BCL-2 inhibition,” Hematology Am. Soc. Hematol. Educ. Program, vol. 2020, no. 1, pp. 1–9, 2020.

[108] S. Qian, Z. Wei, W. Yang, J. Huang, Y. Yang, and J. Wang, “The role of BCL-2 family proteins in regulating apoptosis and cancer therapy,” Front. Oncol., vol. 12, 2022.

[109] https://www.thermofisher.com/us/en/home/life-science/antibodies/antibodies-learning-center/antibodies-resource-library/cell-signaling-pathways/akt-signaling-pathway.

[110] Wikipedia contributors, “AKT1,” Wikipedia, The Free Encyclopedia, 29-Sep-2022. [Online]. Available: https://en.wikipedia.org/w/index.php?title=AKT1&oldid=1112988871.

[111] Genecards.org. [Online]. Available: https://www.genecards.org/cgi-bin/carddisp.pl?gene=AKT1. [Accessed: 10-Dec-2023].

[112] Wikipedia contributors, “Akt/PKB signaling pathway,” Wikipedia, The Free Encyclopedia, 01-Dec-2023. [Online]. Available: https://en.wikipedia.org/w/index.php?title=Akt/PKB_signaling_pathway&oldid=1187729549.

[113] P. Syndrome, “Health conditions related to genetic changes,” Medlineplus.gov. [Online]. Available: https://medlineplus.gov/download/genetics/gene/akt1.pdf. [Accessed: 10-Dec-2023].

[114] P. Syndrome, “Health conditions related to genetic changes,” Medlineplus.gov. [Online]. Available: https://medlineplus.gov/download/genetics/gene/akt1.pdf. [Accessed: 10-Dec-2023].

[115] Wikipedia contributors, “Cyclin-dependent kinase 6,” Wikipedia, The Free Encyclopedia, 07-Nov-2023. [Online]. Available: https://en.wikipedia.org/w/index.php?title=Cyclin-dependent_kinase_6&oldid=1184029332.

[116] L. Zhang et al., “CDK6-PI3K signaling axis is an efficient target for attenuating ABCB1/P-gp mediated multi-drug resistance (MDR) in cancer cells,” Mol. Cancer, vol. 21, no. 1, 2022.

[117] T. Adon, D. Shanmugarajan, and H. Y. Kumar, “CDK4/6 inhibitors: a brief overview and prospective research directions,” RSC Adv., vol. 11, no. 47, pp. 29227–29246, 2021.

[118] L. Zhang et al., “CDK6-PI3K signaling axis is an efficient target for attenuating ABCB1/P-gp mediated multi-drug resistance (MDR) in cancer cells,” Mol. Cancer, vol. 21, no. 1, 2022.

[119] Wikipedia contributors, “Cyclin-dependent kinase 6,” Wikipedia, The Free Encyclopedia, 07-Nov-2023. [Online]. Available: https://en.wikipedia.org/w/index.php?title=Cyclin-dependent_kinase_6&oldid=1184029332.

[120] T. Adon, D. Shanmugarajan, and H. Y. Kumar, “CDK4/6 inhibitors: a brief overview and prospective research directions,” RSC Adv., vol. 11, no. 47, pp. 29227–29246, 2021.

[121] J. P. Alao, “The regulation of cyclin D1 degradation: roles in cancer development and the potential for therapeutic invention,” Mol. Cancer, vol. 6, no. 1, p. 24, 2007.

[122] Sciencedirect.com. [Online]. Available: https://www.sciencedirect.com/science/article/pii/S0167488914001600. [Accessed: 10-Dec-2023].

[123] F. I. Montalto and F. De Amicis, “Cyclin D1 in cancer: A molecular connection for cell cycle control, adhesion and invasion in tumor and stroma,” Cells, vol. 9, no. 12, p. 2648, 2020.

[124] Sciencedirect.com. [Online]. Available: https://www.sciencedirect.com/science/article/pii/S1535610811001780. [Accessed: 10-Dec-2023].

[125] Nature.com. [Online]. Available: https://www.nature.com/articles/sj.onc.1208957. [Accessed: 10-Dec-2023].

[126] C. Busch, T. Mulholland, M. Zagnoni, M. Dalby, C. Berry, and H. Wheadon, “Overcoming BCR::ABL1 dependent and independent survival mechanisms in chronic myeloid leukaemia using a multi-kinase targeting approach,” Cell Commun. Signal., vol. 21, no. 1, 2023.

[127] S. Kamarajugadda et al., “Cyclin D1 represses peroxisome proliferator-activated receptor alpha and inhibits fatty acid oxidation,” Oncotarget, vol. 7, no. 30, pp. 47674–47686, 2016.

